# A new subclass of exoribonuclease resistant RNA found in multiple *Flaviviridae* genera

**DOI:** 10.1101/2020.06.26.172668

**Authors:** Matthew J. Szucs, Parker J. Nichols, Rachel A. Jones, Quentin Vicens, Jeffrey S. Kieft

## Abstract

Viruses have developed innovative strategies to exploit the cellular machinery and overcome the host antiviral defenses, often using specifically structured RNA elements. Examples are found in flaviviruses; during flaviviral infection, pathogenic subgenomic flaviviral RNAs (sfRNAs) accumulate in the cell. These sfRNAs are formed when a host cell 5’ to 3’ exoribonuclease degrades the viral genomic RNA but is blocked by an exoribonuclease resistant RNA structure (xrRNA) located in the viral genome’s 3’untranslated region (UTR). Although known to exist in several *Flaviviridae* genera the full distribution and diversity of xRNAs in this virus family was unknown. Using the recent high-resolution structure of an xrRNA from the divergent flavivirus Tamana bat virus (TABV) as a reference, we used bioinformatic searches to identify xrRNA in the *Pegivirus, Pestivirus, and Hepacivirus* genera. We biochemically and structurally characterized several examples, determining that they are genuine xrRNAs with a conserved fold. These new xrRNAs look superficially similar to the previously described xrRNAs but possess structural differences making them distinct from previous classes of xrRNAs. Our findings thus require adjustments of previous xrRNA classification schemes and expand on the previously known distribution of the xrRNA in *Flaviviridae*, indicating their widespread distribution and illustrating their importance.

**IMPORTANCE:** The *Flaviviridae* comprise one of the largest families of positive sense single stranded (+ssRNA) and it is divided into the *Flavivirus*, *Pestivirus*, *Pegivirus*, and *Hepacivirus* genera. The genus *Flavivirus* contains many medically relevant viruses such as Zika Virus, Dengue Virus, and Powassan Virus. In these, a part of the virus’s RNA twists up into a very special three-dimensional shape called an xrRNA that blocks the ability of the cell to “chew up” the viral RNA. Hence, part of the virus’ RNA remains intact, and this protected part is important for viral infection. This was known to occur in Flaviviruses but whether it existed in the other members of the family was not known. In this study, we not only identified a new subclass of xrRNA found in Flavivirus but also in the remaining three genera. The fact that this process of viral RNA maturation exists throughout the entire *Flaviviridae* family makes it clear that this is an important but underappreciated part of the infection strategy of these diverse human pathogens.

## INTRODUCTION

Viruses face continuous evolutionary pressure to evolve innovative strategies that exploit the host cell′s biological machinery and overcome its antiviral defenses. Often, these are based on specifically structured RNA elements, which is not surprising given RNA′s functional diversity and ability to influence virtually every cellular process. Important examples are found in positive sense single stranded RNA (+ssRNA) viruses such as arthropod-borne flaviviruses, which use several structured elements within their RNA genome to direct or regulate important processes during infection. (1–13).

Among the important structured RNA elements found in flaviviruses are exoribonuclease-resistant RNAs (xrRNAs), which enable an elegant mechanism of non-coding RNA biogenesis. Originally identified in mosquito borne flaviviruses (MBFVs) (14–19) and later in +ssRNA plant-infecting viruses (20, 21), xrRNAs block the processive degradation of the viral genome by host cell 5′-3′ exoribonuclease Xrn1(19), the enzyme responsible for the majority of cytoplasmic RNA decay (Fig. 1A) (22). During infection, a subset of flaviviral genomes is targeted to the cellular RNA decay machinery, then processively degraded in a 5′ to 3′ direction by Xrn1, until the enzyme halts at an xrRNA in the 3′ untranslated region (UTR) (19, 23). The fold of the xrRNA is sufficient for this function, and no accessory proteins nor chemical modification of the RNA are required. The xrRNA and the protected downstream RNA comprise a non-coding subgenomic flaviviral RNA (sfRNA) that accumulates in the cell and performs several important functions for the virus (2, 19, 23–26), including inhibiting host antiviral response (27, 28), affecting viral transmissibility (29–33), and disrupting the host mRNA program (34, 35). Within plant viruses, xrRNAs are associated with both non-coding and coding subgenomic RNAs (36).

**Fig. 1.**
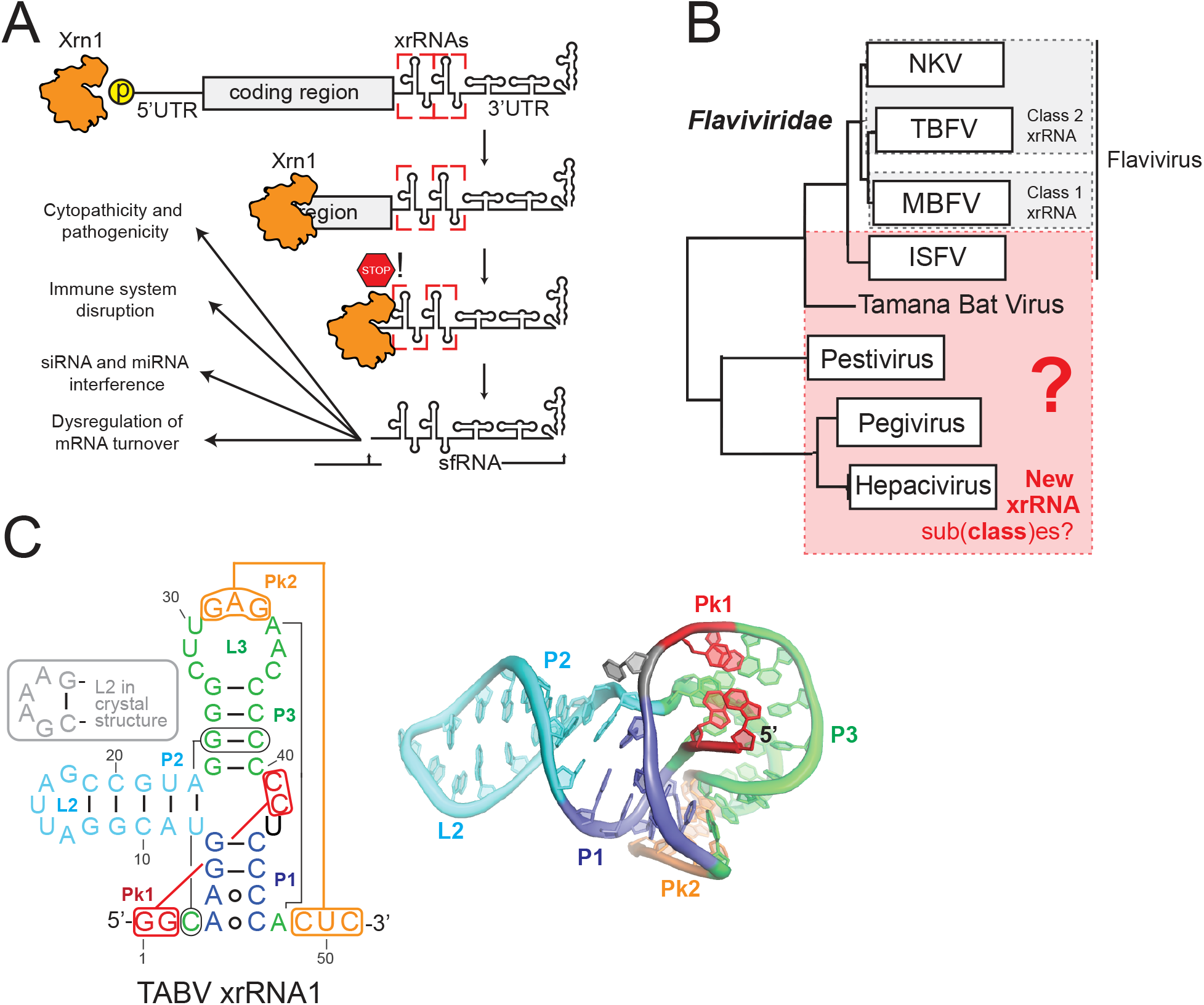
Mechanism and presence of exoribonuclease resistant RNA in Flaviviridae. (A) Mechanism of sfRNA biogenesis through Xrn1 exoribonuclease (orange) resistant RNA (xrRNA, boxed in red). (B) Phylogeny of *Flaviviridae* based on the sequence of NS5 adapted from (44). Genera possessing class 1 or class 2 xrRNAs are boxed in grey. Genera investigated in this study are boxed in red. NKV, no-known vector flavivirus; TBFV, tick borne flavivirus; MBFV, mosquito-borne flavivirus. (C) Secondary structure diagram of the xrRNA1 from TABV. Secondary structure regions are delineated by different colors and key long-range interactions are indicated with lines. (D) Three-dimensional structure of TABV xrRNA colored as in panel C. (PDB ID XXXX) (37).

Detailed three-dimensional crystal structures of xrRNAs from several flaviviruses provide insight into how an xrRNA resists the action of 5′-3′ exoribonucleases (1, 13, 37). Specifically, all xrRNAs fold into a compact structure containing a ring-like feature that wraps around the 5′ end of the resistant RNA element. Biochemical analyses suggest that this topology acts as a molecular brace against the surface of the exoribonuclease approaching from the 5’ side, preventing it from progressing past a defined point (38). This ring is formed through specific structural motifs, including a three-way junction, pseudoknots and other long-range tertiary interactions, non-Watson-Crick base pairs, complex base stacking arrangements, and base triples (39). A ring-like structure is also found in the xrRNAs from some genera of the plant-infecting *Tombusviridae* and *Luteoviridae* viral families although the structural strategy used to form and stabilize the ring is different (21, 36, 40). There may be yet-undiscovered ways to form a ring-like structure and thus other RNA architectures that can block exoribonucleases.

Biochemical, structural, and sequence comparison studies led us to group flaviviral xrRNAs into two distinct classes (Fig. 1B) (38). The class 1 xrRNAs are found in the MBFV, some no known vector (NKV) flaviviruses, and some insect-specific flaviviruses (ISFV). The class 2 xrRNAs are found in tick-borne flaviviruses (TBFV) and some related NKV viruses. These classifications are based on the proposed secondary structure of the elements, the halt-point of Xrn1 relative to these putative secondary structures, and sequence conservation patterns (38). For example, in the class 1 the sequence of the P1 stem is highly conserved, but the same sequences are not found in the class 2. Classification is informed and refined by three-dimensional structures which improve sequence alignment efforts [for class 1 MBFV xrRNAs (1, 13), and for an xrRNA from Tamana bat virus (TABV)(37); there is no structure of a class 2 yet].

The *Flaviviridae* family consists of four major genera: *Flavivirus*, *Pegivirus*, *Pestivirus*, and *Hepacivirus* (41–44). Within *Flaviviridae*, xrRNAs have been definitively identified only in the *Flavivirus* genus, in a single pestivirus (which might actually be a flavivirus) (3, 45, 46) and now the phylogenetically isolated Tamana bat virus (Fig. 1B)(37). If xrRNAs exist in the genomic RNA of the *Pegivirus*, *Pestivirus*, and *Hepacivirus* genera, this would indicate their presence in the three *Flaviviridae* genera that were previously thought to be devoid of xrRNAs. Their fold may also look different, therefore potentially calling for classification adjustments as more viruses are discovered, more xrRNAs are characterized, and more detailed structural information is gained.

New questions about variation in flavivirus xrRNA structure arise from the recent structure of an xrRNA found in the 3′ UTR of TABV (37). While this xrRNA superficially resembles a class 1 xrRNA, it lacks the conserved sequences essential to form the required tertiary contacts in the class 1 (Fig. 1C)(38). Consistent with this, although the TABV xrRNA structure forms the ring-like fold, it uses an unexpected set of tertiary interactions compared to the MBFV xrRNA (Fig. 1C) (37). This divergence is different enough to suggest a potentially new xrRNA subclass with other unidentified members, and to raise the question of whether additional classes of xrRNAs exist among the *Flaviviridae*.

Using computational tools that take into account both structural and sequence constraints, and informed by crystal structures, we exploited the burgeoning availability of genomic data from new viral species. We identified putative xrRNAs in the *Pegivirus*, *Pestivirus*, and *Hepacivirus* genera that we verified using *in vitro* functional studies and characterized by chemical probing. Analysis of sequences and secondary structures revealed fundamental similarities to the TABV xrRNA. Together, these xrRNAs comprise an xrRNA subclass similar to the class 1 but bioinformatically and structurally distinct; we now assign them to subclass 1b and accordingly place the previously classified species in class 1 (MBFV) into subclass 1a. Thus, our results show that all genera (although not necessarily all species) within *Flaviviridae* contain xrRNAs that exist in at least three distinct structural classes/subclasses.

## RESULTS

### Computational identification of an xrRNA subclass

The recent crystal structure of an xrRNA from TABV (37) revealed a previously unknown secondary structure, which motivated reexamination of existing alignments and secondary structures of many putative xrRNAs from insect specific flaviviruses (ISFV). Therefore, we constructed a new initial sequence alignment from 20 likely xrRNA sequences belonging to ISFV, taking into account the patterns revealed in the TABV structure. These sequences often occur in multiple copies in series, and thus we included all of these in our alignment (referred to as xr1, xr2, etc. in the 3′ UTR of CxFV, QBV, MOsFV, CxThFV, KRV, AeFV, CFAV; see Table S1 and S2 in the supplemental material) (47). These ISFV xrRNA sequences had previously been proposed to conform to class 1 xrRNA or to have different secondary structures (38, 47), but we could align them more convincingly to the TABV xrRNA, using information from its crystal structure (37) (see Methods). The resulting initial covariance model was used as input to the program Infernal to search for a similar structure pattern through all available +ssRNA viral sequences at the National Center for Biotechnology Information (NCBI; last retrieved on 04/24/2020) (48). Multiple iterations (see Materials and Methods) resulted in the identification of 66 putative xrRNA sequences, belonging to the following genera: *Flaviviruses* (n = 23), *Pegiviruses* (n = 27), *Pestiviruses* (n = 7), and *Hepaciviruses* (n = 9) (Fig. 2A; see also Table S1 in the supplemental material). Not all known viral species had xrRNAs that could be identified by our bioinformatic searches, as many sequences in the database did not include the 3′ UTR. As such, more viruses may contain putative xrRNAs, which were not identified here due to incomplete reporting of the viral genome sequences.

**Fig. 2:**
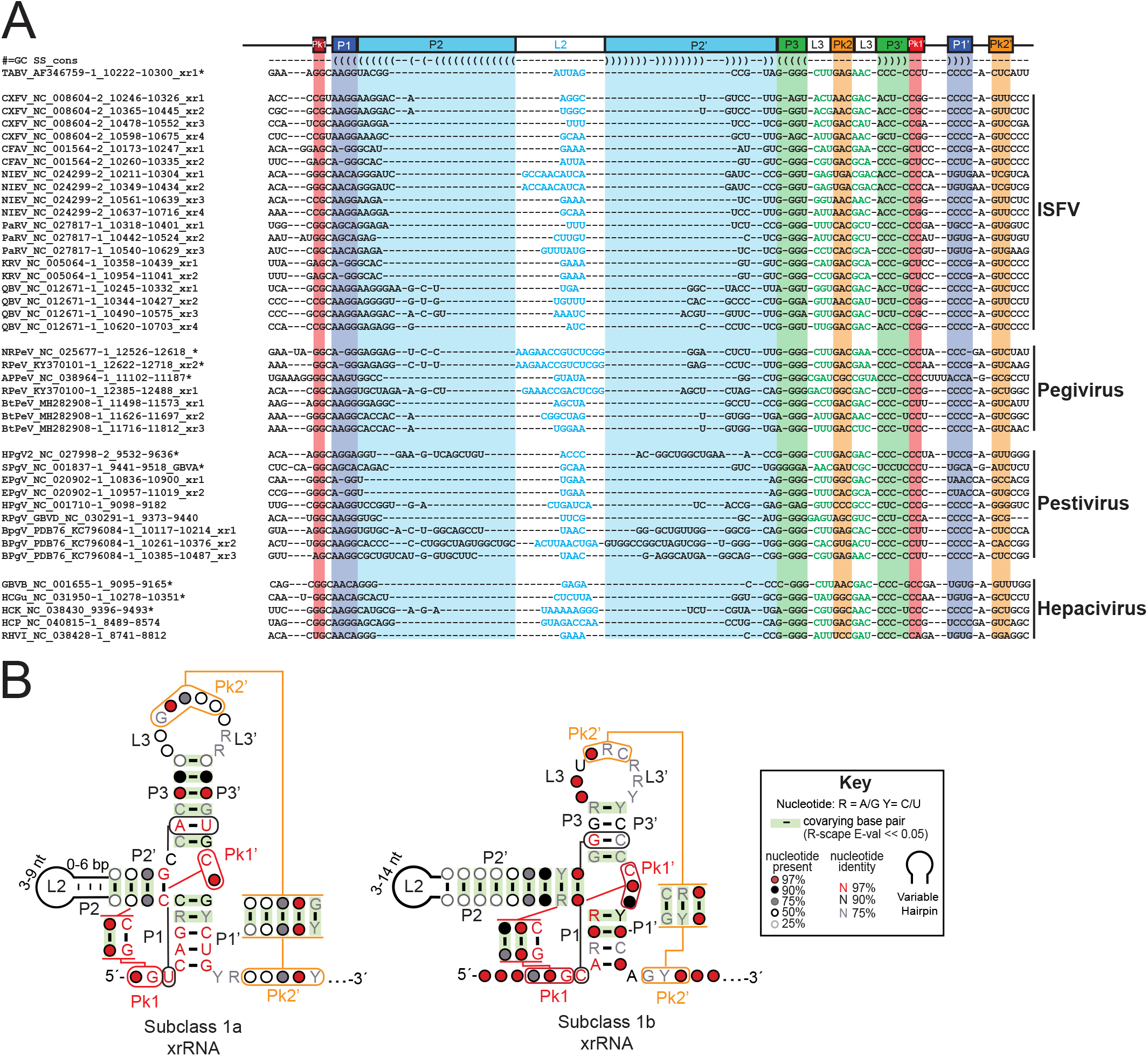
Sequence conservation of subclass 1b xrRNA. (A) Comparative sequence alignment of select subclass 1b xrRNA from insect specific flavivirus (ISFV), *Pestivirus, Pegivirus*, and *Hepacivirus*. The TABV xrRNA xr1 sequence and the secondary structure derived from the crystal structure were used as a reference. A star (*) designates sequences used for subsequent biochemical analysis. (B) Covariance models of the subclass 1a (32 sequences; updated from (39)) and subclass 1b (87 sequences) xrRNAs. Covarying base pairs validated by R-scape (E-value < 0.05) are highlighted in green.

From this expanded comparative sequence alignment, we generated a covariance model of the newly proposed xrRNAs (Fig. 2B, right). The comparative sequence alignment, covariance model, and the fact that these xrRNA are bioinformatically distinct from the MBFV, reveals that these identified and revised xrRNA’s did not fit into the class 1 xrRNA as it was currently defined. Thus, we divided the current class 1 xrRNA into two subclasses: subclass 1a (previously defined class 1 xrRNA) and subclass 1b (this current work). Differences observed between the subclass 1b and the subclass 1a were the sequence patterns of their P1 and P3 stems, nucleotide patterns of the L3 loop, Pk1 region, and the presence or absence of a nucleotide between P2 and P3. These are analyzed and discussed in more depth below.

Analysis of the covariance model was performed with R-scape (49) which gave statistical support to the model and the proposed three stems (P1, P2, & P3). Specifically, in each stem, E-values for covarying base pairs ranged from 8.89 × 10^−7^ – 1.29 × 10^−5^ (P1), 2.17 × 10^−14^–1.75 × 10^−3^ (P2), and 1.45 × 10^−4^–1.77x 10^−9^ (P3). Stem length varies less for P1 (3–5 base pairs) and P3 (4–5 base pairs), than for P2 (up to 21 predicted base pairs) (Fig. 2B). This variation in stem length had been previously noticed for class 1 MBFVs, albeit with different stem length requirements (P1, 5 base pairs; P3, 4–8 base pairs; but P2, 1–9 base pairs). The similarly conserved lengths of P1 and P3 can be explained by their participation in ring formation, while P2 extends away from the core fold, explaining its variable length (Fig. 1C) (37). The anticipated Pk1 and Pk2 pseudoknots that we manually predicted based on the TABV structure were also supported by R-scape (Pk1 E-values: 4.58×10^−8−^ 1.20×10^−13^) (Fig. 2B).

All flavivirus class 1 xrRNAs contain a base triple necessary for forming the functional fold. In the MBFV this is a U…A-U triple while in TABV it is replaced by an isosteric C…G-C (37). In our covariance model, sequence conservation of this C…G-C base triple is >97% for G, and >97% and >90% for the two Cs (Fig. 2B). Only two sequences identified in our alignment (CXFV_NC_008604.2 Xr1 & Xr4) possess a U…A-U instead of the C…G-C indicating an instance of covariation but not enough to be supported by R-scape (Fig. 2A; see also Table S1 in the supplemental material). Finally, none (except one) of the sequences in our alignment contain any nucleotide between P2 and P3, a peculiarity that was discussed in the report of the TABV xrRNA structure (37). The one sequence that does contain a nucleotide at that position comes from *Rodent hepacivirus* isolate rn-1 (RHV; Table S1). This putative xrRNA has a U in this position, which cannot be otherwise accommodated within P2 or P3. Similarly, other *Hepacivirus* isolates belonging to sequences such as the *Rodent hepacivirus* isolate RrMC-HCV (RtMC, Table S1) comprise sequences that depart from the subclass 1b pattern, potentially leading to non-Watson-Crick pairs at the base of P2 or within a somewhat altered three-way junction (sequences marked by ′!!′ in Table S1). These sequences are rare and found mainly in members of the *Hepacivirus* genera.

Another subset of sequences deviates from the covariance model at the base of P1. They comprise putative ISFV xrRNA sequences that are found in the more downstream copies of putative xrRNA (xr3, xr4, etc.). Among them are xr4 from the *Parramatta River virus* isolate 92-B115745 xr4 (PARV_xr4; Table S1), xr3 from Cell fusing agent virus strain Galveston (CFAV_xr3; Table S1), xr1 from Mercadeo virus isolate ER-M10 (MECDV_xr1; Table S1) and xr1 from Menghai flavivirus isolate MHAedFV1 (MFV_xr1; Table S1). For example, only a two base-pair P1 can be proposed for PaRV_xr4, with no equivalent to A48 in TABV. Further testing will be required to determine whether these sequences indeed correspond to xrRNAs, and if so, whether they possibly form a subtype within subclass 1b.

### Bioinformatically identified subclass 1b xrRNAs are exoribonuclease resistant

To test whether computationally identified subclass 1b xrRNAs were resistant to Xrn1 and therefore comprised authentic xrRNAs, we subjected representative sequences from *Pestiviruses, Pegiviruses and Hepaciviruses* to *in vitro* Xrn1 resistance assays (Fig. 3). Briefly, eight putative subclass 1b xrRNAs (marked by * in Fig. 2A) were transcribed *in vitro* with a 22 nt leader sequence, which allows Xrn1 to load onto the 5′ end of the transcribed RNA. Some of the previous work from our laboratory showed that a wild-type leader sequence of an arbitrary length could generate false negative results in Xrn1 resistance assays, likely due to RNA misfolding. Hence, for our assays of subclass 1b xrRNAs, the leader sequence contained the same normalization hairpin (xxxx-GAGUA-xxxx) and spacer sequences used for the chemical probing experiments described in the next section (Methods; see Table S3 in the supplemental material). This adjustment also contributed to consistency between resistance assays and probing experiments.

**Figure 3:**
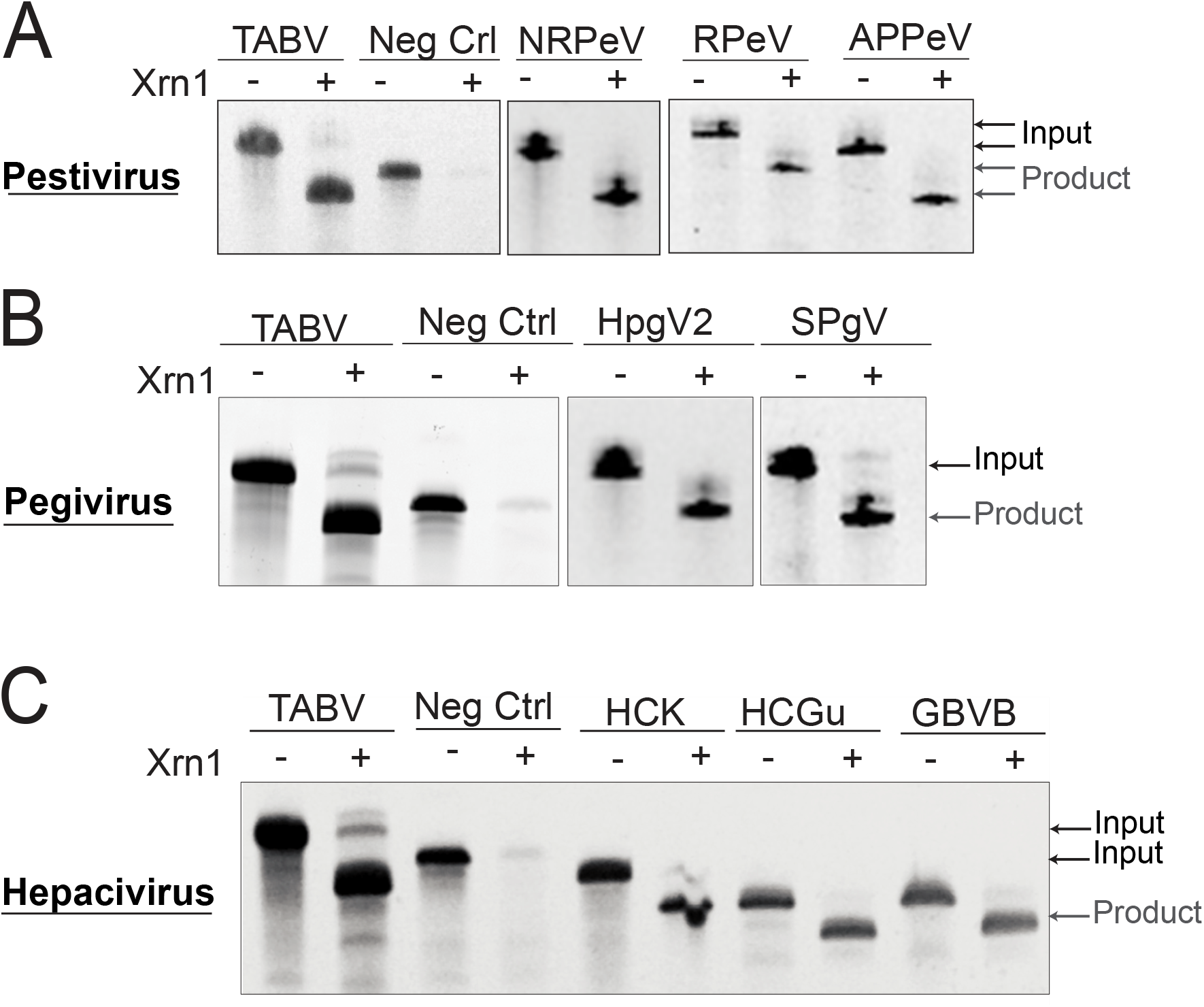
Biochemical validation of exoribonuclease resistance of subclass 1b xrRNA representatives. Each panel shows the results from denaturing 8% polyacrylamide gel electrophoresis used to analyze Xrn1 resistance experiments. (A) Representative Pestivirus subclass 1b xrRNA. From left to right: NRPeV, Norway Rat Pestivirus (NRPeV_NC025677-1_12526-12618*); RPeV, Rodent Pestivirus (RPeV_KY370101-1_2622-12718_xr1*); APPeV, Atypical Porcine Pestivirus (APPeV_NC_038964-1_11102-11187). (B) Representative subclass 1b xrRNA from Pegivirus. From left to right: Simian Pegivirus (SPgV_NC_001837-1_9441-9518_GBVA*) and Human Pegivirus 2 (HPgV2_NC_027998-2_9532-9636_CP*). (C) Representative subclass 1b xrRNA from Hepacivirus. From left to right: GB virus B (GBVB_NC_001655-1_9095-9165*), Guareza Hepacivirus (HCGu_NC_031950-1_10278-10351) Hepacivirus (HCGu_NC_031950-1_10278-10351*). TABV, Tamana bat virus (positive control); Neg Ctrl, non-resistant RNA (tRNA-like structure) (51).

RNAs were treated with RNA 5′-Pyrophosphohydrolase (RPPH), leaving a mono-phosphorylated 5′ end (50). When an RNA is resistant to Xrn1, subsequent incubation with recombinant Xrn1 leads to a defined but smaller size product when resolved by polyacrylamide electrophoresis, as the enzyme has loaded, partially degraded the RNA, but then stopped. As a positive control, we used the *in vitro* transcribed TABV xrRNA (xr1) (Table S3). An RNA with a tRNA-like structure that is not resistant to Xrn1, with appended normalization hairpins, was used as a negative control (51). When challenged with Xrn1, all putative xrRNA sequences identified in our computational searches were resistant, indicated by the appearance of the shorter but stable RNA product (Fig. 3). This result confirmed that even sequences with very short (3 nt) (Fig. 3C: GBVB) or long P2 stems (15 and 10) (Fig. 3B: HPgV2, Fig. 3C: HCK), or with the potential for a particularly long Pk1 (4nt) (Fig. 3B: SPgV) can form an Xrn1 resistant structure. Even the xrRNA from HCP, which displays a putative A.A pair at the base of P2, is resistant to Xrn1 (see Fig. S1 in the supplemental material). Thus, the bioinformatically identified subclass 1b xrRNAs that we tested are authentic exoribonuclease-resistant elements and this result suggests that the untested sequences from our sequence alignment likely are also true xrRNAs.

### Secondary structure validation and chemical probing of subclass 1b xrRNAs

Since the secondary structures proposed on the basis of sequence alignments and covariation are predictions, we tested them experimentally through chemical probing with dimethyl sulfate (DMS; modifies the Watson-Crick side of unpaired A and C) and N-methylisatoic anhydride (NMIA; modifies the 2′ hydroxyl group of nucleotides not involved in Watson-Crick pairs) (Fig. 4) (52, 53).

**Figure 4:**
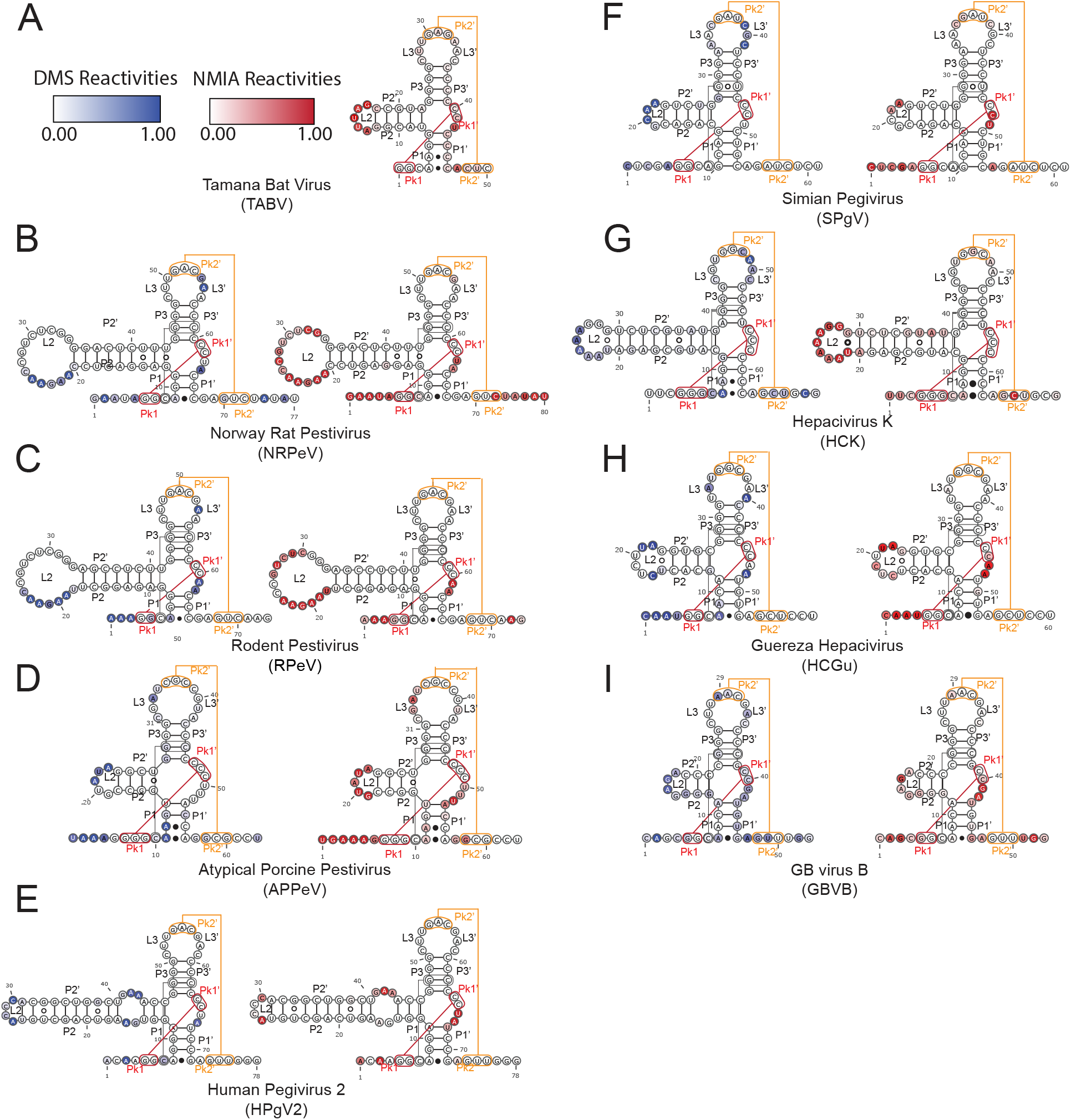
Secondary structure validation of the subclass 1b through chemical probing. Subclass1b constructs were probed with DMS and NMIA. Normalized DMS (blue) and NMIA (red) reactivities were combined with comparative sequence analysis to derive secondary structure models. Reactivities were overlaid on the secondary structure using Varna v. 3.93 (64). (A) Tamana Bat Virus. (B–D) Pestivirus. (E–F) Pegivirus. (G–I) Hepacivirus. Raw and normalized data shown in Fig. S2 of the supplemental material.

We first probed (using NMIA only) the wild-type sequence of the TABV xrRNA, which expands on observations based on the crystal structure by providing a ′fingerprint′ of the chemical probing pattern for this type of fold (Fig. 4A). Most of the nucleotides were unreactive to NMIA (including in L3 which appears as ′unpaired′ in the secondary structure diagram, except for Pk2), supporting a compact and stable fold. Highly reactive positions in the exposed loop L2 were consistent with the TABV xrRNA crystal structure (Fig. 1C). Likewise, strong reactivities at U43 and A48 can also be rationalized by the structure: U43 is flexible and the ribose of A48 adopts a reactive C2′ endo pucker (54, 55). Although chemical probing was performed with the wild-type TABV sequence, the resulting data are consistent with the structure of the sequence variant engineered for crystallization (37). In previous studied, removal of the P4 stem has did not affect Xrn1 resistance (37, 56), thus we did not examine it as part of our analysis but the full chemical probing construct with the P4 stem is displayed in the supplementary information (Fig. S2)

We performed chemical probing of the eight *Pegivirus, Pestivirus*, and *Hepacivirus* xrRNAs tested for Xrn1 resistance to compare the probing ‘fingerprint’ to that of TABV. The resultant patterns are consistent with the pattern from TABV xrRNA, indicating that they adopt a similar secondary structure (Fig. 4). In seven cases, the RNAs were similarly non-reactive overall, except for L2 and the nucleotide equivalent to U43 in TABV (Fig. 4B–H). GBVB showed the same pattern but was overall more reactive, likely indicative of a less stable *in vitro* fold (Fig. 4I). Several of the RNAs contained >30 nucleotide expansions in the P2 regions, and in all cases the probing indicated they were folded as elongated stem loops, in some cases with internal loops as seen in HPgV2 (Fig. 4B,C,E,G). In some of these, parts of L2 may be involved in additional intramolecular interactions, as suggested by the absence of reactive positions within L2 for NRPeV, RPeV and HPgV2 (Fig. 4B, C, E). Probing revealed that some sequences had two (NRPeV, RPeV, HPgV2, GBVB) or even three (APPeV) reactive nucleotides between Pk1′ and P1′, regardless of the potential for Pk1 to contain more than three base pairs, which could be interpreted from only looking at the sequence (Fig. 4B–E, I). Only one sequence tested (HCK) had no reactive nucleotide at that position (Fig. 4G).

Intramolecular interactions involved in proper ring folding, such as the equivalent position to the strongly reactive A48 in TABV, were reactive only in APPeV (G57), SPgV (A54) and GBVB (A47) (Fig. 4D,F,I). Finally, reactive positions to DMS but not NMIA in L3′ (the 3′ side of L3) for NRPeV, RPeV, SPgV, HCK, HCGu, and GBVB implied the Watson-Crick face of these residues remained available in an otherwise structured region of stacked bases, which is compatible to the availability of the A34, A35, and C36 in the the TABV crystal structure. Overall, the chemical probing data support the predicted secondary structure models that indicate these xrRNAs share the same TABV xrRNA-like fold, consistent with their inclusion in subclass 1b.

### Structural analysis of the subclass 1b xrRNA

Based on the sequence covariation observed and our structural analysis, xrRNAs from subclass 1b are more compact than those from subclass 1a, except for the P2 stem (Fig. 2B). Subclass 1b xrRNAs are distinguished from subclass 1a by four structural features observed in the TABV xrRNA crystal structure. The first is the presence of generally two non-Watson-Crick interactions at the 5′ end of the P1 stem (Fig. 5A, 5C). Over half of the sequences in our alignment have the two A…C interactions observed in the TABV structure, followed by two Watson-Crick pairs before the three-way junction (Fig. 2A,; see also Table S1 in the supplemental material). The interactions between A4…C46 and A5…C45 seen in the TABV structure help create a sharp bend in the structure associated with a break in base stacking between Pk1 and P2 (Fig. 5A). For the remainder of the sequences, one of the two As is missing, probably the equivalent to A5, because A4 is involved in stabilizing interactions at the ring closure (Fig. 5A). In 28% of sequences, A4…C46 is replaced by A…A/G interactions. This change is accompanied by the presence of A–U or G–C instead of A5…C45, suggesting a third Watson-Crick pair could form in P1 in that case. A common Hoogsteen/sugar edge configuration at A…G and Watson-Crick at A–U or G–C would lead to a shift of the pairing partner of both A bases towards the minor groove. In any event, the length of P1 is constrained to 3–4 base pairs by the compactness of the fold, a structural feature which is supported by our comparative sequence alignment (Fig. 2A). Conversely, the subclass 1a P1 stem is highly conserved with only two base pairs covarying out of the six possible Watson-Crick base pairs. Also, the subclass 1b lack almost absolutely-conserved sequences that are found in the subclass 1a xrRNAs, for example in the P1 stem, making them distinct (39).

**Figure 5:**
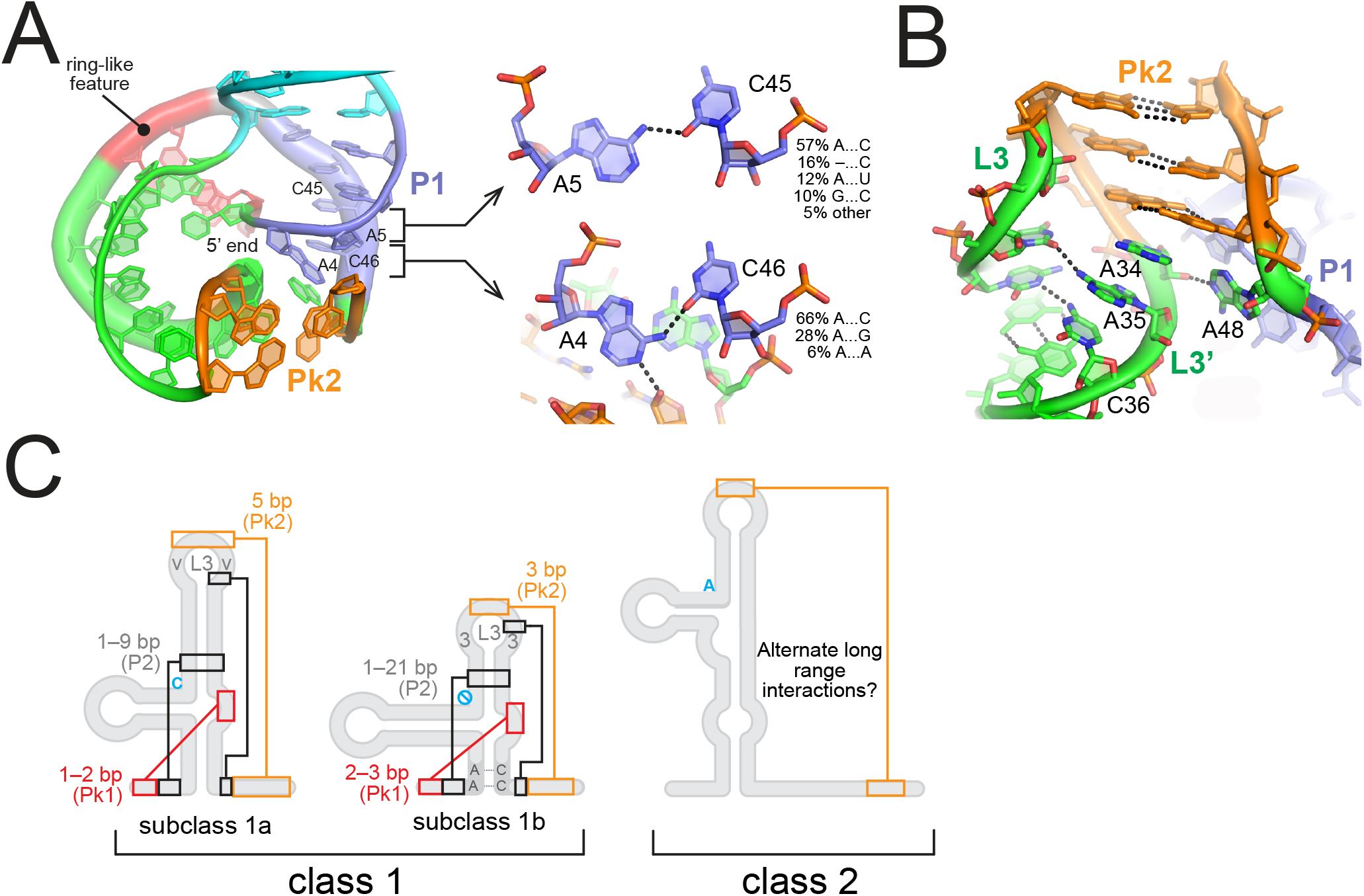
Structural analysis of the subclass 1b xrRNA. (A) View of the ring-like structure in the TABV xrRNA from the 3′ side. The 5′ end is kept at the center of the ring and points away from the reader. Regions are color-coded as in Fig. 1C. A…C interactions in P1 are highlighted in the close ups. (B) Close up of L3 and Pk2. (C) Structural features for classes 1 and 2 are highlighted on secondary structure silhouettes. The blue “no” symbol for subclass 1b highlights the absence of a nucleotide in the junction between P2 and P3. In addition to the differences highlighted here, subclass 1a contains regions of absolutely-conserved sequences that are not in subclass 1b (Fig. 2B)

The L3 loop comprising the Pk2 pseudoknot is the second defining feature of the subclass 1b xrRNA. The L3 region exists in a 3-3-3 (86%), 4-3-3/3-3-4 (13%), or a 4-3-4 (1%) nucleotide configuration (the number of nucleotides refers to the length of the 5′ side of L3, of Pk2, and of the 3′ side of L3 or L3′; Fig. 2A, 5C, Table S1). Sequence patterns of L3 can be rationalized based on the TABV structure: (i) the 5′ side generally comprises three pyrimidines (90% U at the position preceding Pk2; Figure 2), due to the tight turn leading to Pk2; (ii) the three base-pair length of Pk2 is conserved as A34 within L3′ immediately downstream participates in ring closure interactions (Figures 2, 5B); (iii) L3′ following Pk2 generally comprises two purines (A34 and A35 in TABV) which help support stacking of Pk2 on P3, followed by a pyrimidine (C36) which could accommodate the guanine present on the 5′ side of L3 in 45% sequences (Figure 2; see also Table S1 in the supplemental material). Notably, the first adenines following Pk2 do not get modified by NMIA, although they are reactive to DMS (Fig. 4), which is consistent with them being conformationally constrained but with available Watson-Crick edges, as revealed by the TABV xrRNA structure (Figure 5B).

Next, a defining structural feature of subclass 1a lacking in subclass 1b is the presence of a conserved cytosine between P2 and P3. In subclass 1a xrRNA, this nucleotide is important for tertiary interactions that support folding and the formation of the ring-like structure around the 5′ end of the xrRNA. Studies have shown that mutating this C to any other nucleotide disrupts the ability of xrRNA to resist Xrn1 (1). In recent studies with TABV it was shown that its fold cannot tolerate the addition of a nucleotide in this region (37). Because the ring of subclass 1b TABV xrRNA is stabilized by a different set of long-range interactions compared to the subclass 1a ZIKV xrRNA, this results in a more compact fold that can no longer accommodate the C. Although this C is critical for subclass 1a xrRNAs, our present work shows that its absence is a shared feature among subclass 1b xrRNAs.

Finally, the covariation pattern of the Pk1 is another key feature of the subclass 1b xrRNA, although less distinctive between the two subclasses. According to sequence alignment and probing data, this region accommodates either two (11% of the sequences identified) or three base pairs (88.3% of the sequences) (Fig. 5B, 5C; see also Table S1 in the supplemental figures). Out of the eight sequences we probed, five reveal that the Pk1 predicted to have three base pairs (based on sequence) actually only forms two base pairs (Fig. 4B, E, F, H, I). Together with the crystal structure of TABV (where Pk1 comprises two base pairs), these results suggest that two base pairs are sufficient to support a resistant fold, although we cannot exclude that Pk1 may have an additional pair on its 5′ side in some other cases, or that this pair is dynamic (Fig. 1C). The second of the two Pk1 base pairs (which is always formed) is >97% a G-C (Fig. 2B), as in subclass 1a (Fig. 2A; reference to Kieft RNA Biology 2015). The first nucleotide immediately downstream of Pk1′ is strongly reactive to NMIA, except for HCK (Fig. 4G). This feature is consistent with the flipped-out conformation of U42 in the TABV structure (Fig. 1B). In the subclass 1a the PK1 region is never more than two base pairs, but it can which can be up to three for the subclass 1b.

## DISCUSSION

In this study we have identified and structurally characterized xrRNAs in all genera of *Flaviviridae*, one of the largest families of RNA viruses, expanding on the known distribution of these RNA structures within *Flaviviridae* (23). Based on our computational, structural, and biochemical data, we propose that the characteristics of these new xrRNAs require a division of the previously proposed class 1 xrRNA (38) into two distinct subclasses: subclass 1a (comprising MBFV, in particular MVEV, ZIKV, WNV, YFV, etc.), and subclass 1b (comprising TABV, ISFV, *pegi-*, *pesti*- and *hepaciviruses*). Subclasses 1a and 1b deviate in (i) the sequence patterns of their P1 and P3 stems, (ii) the nucleotide patterns of the L3 loop, (iii) the Pk2 region, and (iv) whether a particular nucleotide is present between P2 and P3. In addition, the pattern of almost absolutely-conserved nucleotides that is indicative of subclass 1a (39), is not present in subclass 1b. Thus, all of the identified xrRNAs in *Flaviviridae* fall into three distinct groups: subclasses 1a and 1b, and class 2 (which remains as described in the literature) (38).

Within known *Flaviviridae* sequences, subclasses 1a and 1b do not appear in the same viral species. In other words, no viral sequence within the phylogenetic groups characterized as having subclass 1a xrRNAs possesses a subclass 1b xrRNA and vice-versa. When several xrRNAs are present in the same 3′ UTR (57), all belong to the same subclass. This observation supports the hypothesis of the evolution of these structural elements from a common ancestor (58), as opposed to horizontal gene transfer. This scenario is further supported by our observation that hepaciviruses — one of the most distantly related genera within *Flaviviridae —* also contain the most divergent subset of subclass 1b xrRNAs, possibly branching out further into distinct subcategories. Moreover, hepaciviruses include viruses like Hepatitis C, which do not have xrRNAs (59). In short, xrRNAs are evolving with the rest of the attached viral genome and not actively ‘hopping’ across genera.

Although subclass 1b xrRNAs are resistant to Xrn1 *in vitro*, their function in the context of viral infection and pathogenesis remains unclear. A key functional role of xrRNAs from subclass 1a is to enable formation of sfRNAs, which have key roles for virus survival within the vector and host cells (23, 27, 60). Whether the subclass 1b xrRNAs from pesti-, pegi- and hepaciviruses have a similar function in the generation of viral sfRNA is currently unknown. In particular, not every virus with a subclass 1b xrRNA generates sfRNAs. As an example, we show that GBV-C has a subclass 1b xrRNA (Fig. S1), although previous studies indicated it may not produce any sfRNA (3). Functions other than generating subgenomic RNAs were also expected from our discovery that plant virus xrRNAs could be located upstream of coding regions (36). Systematically testing of viruses for formation of sfRNAs would be a first step in pinpointing alternative functions of subclass 1b xrRNA. Whether subclass 1b xrRNAs participate in sfRNA generation or not, evolutionarily they maintain the necessary interactions for proper folding into a structure that can resist exoribonucleases.

In addition to motivating systematic characterization of sfRNA formation within all *Flaviviridae* genera, this study also reinforces the importance of combining bioinformatic, biochemical, and structural techniques to derive meaningful conclusions about viral structured RNAs. Bioinformatics are a powerful tool to develop xrRNA secondary structure predictions, but this approach requires the availability of complete 3′ UTR sequences. We estimate that >50% of the available viral genome sequences for Pesti-, Pegi- and Hepaciviruses end at the last codon of the coding region. In addition, biochemical experiments are needed to not only validate computational predictions, but also to refine the proposed models. As an example, the APPeV xrRNA was predicted to have a six-base pair Pk1, based solely on the sequence alignment and our covariance model. This possibility was refuted — in the context of Xrn1 resistance at least — after the chemical probing experiments revealed that Pk1 consisted of three base pairs (Fig. 4D). We also bioinformatically identified two putative xrRNAs in HPgV2 whose sequences were overlapping (Fig. S2). The Xrn1 resistance assay enabled us to determine which sequence corresponded to the true xrRNA. Similarly, further testing and structural mapping of the putative but still ambiguous xr3–5 elements for some of the ISFVs (CSFV and PaRV) should reveal whether they are actual xrRNAs, and if so, whether they might represent another subgroup within subclass 1b. Overall, this study expands on the widespread presence of xrRNAs in nature, now within *all Flaviviridae* viruses, thereby further supporting the prevalence of this particular structure in the viral world.

## METHODS

### Subclass 1b bioinformatic searches and covariance model analysis

An initial subclass 1b alignment was created starting from a total of 20 sequences of insect specific flaviviruses (see Table S1, sequences with a ‘+’ symbol) that were manually aligned with a combination of *RALEE* v. 0.8 and a text editor, using the TABV secondary structure as a reference. Using *Infernal* v. 1.1.3 (48) with default parameters we searched a database of reference viral genomes consisting of all currently available +ssRNA sequences downloaded from the National Center for Biotechnology Information (NCBI; last retrieved on 04/24/2020). Hits from the Infernal searches were manually added to the comparative sequence alignment, when they met all of the following criteria: *Infernal* E value <0.05, >15% nucleotide variation within each sequence, presence of Pk1 and Pk2, location within the 3′ UTR. Sequences with higher E values were also inspected and added to the list if they met the last three requirements. For the final proposed covariance model of 87 xrRNA sequences, we performed statistical validation using *R-scape* v.1.5.3 (49) and rendering with *R2R* v.1.0.5 (61).

### Plasmid construction

Each xrRNA construct (Table S3) was designed as a double stranded DNA gBlock (IDT) and was subsequently cloned into the EcoRI and BamHI sites of pUC-19. Cloned plasmids were amplified in competent *E. coli* DH5α cells and purified with Qiagen miniprep kit (Qiagen). The recovered plasmid stocks were verified through sequencing (Eton Bioscience).

### In vitro RNA transcription

Template DNA was amplified by PCR using custom DNA primers (Table S3) and recombinant Phusion Hot Start polymerase (New England Biolabs). *In vitro* transcription was carried out in a volume of 2.5 mL comprising 1.0 mL of PCR reaction as template. The transcription reaction contained ~0.2 M template DNA, 6 mM each rNTP, 60 mM MgCl_2_, 30 mM Tris pH 8.0, 10 mM dithiothreitol (DTT), 0.1% spermidine, 0.1% Triton X-100, T7 RNA polymerase, and 2 μL RNasin RNase inhibitor (Promega). The transcription reaction was incubated overnight at 37°C. The RNA was precipitated with 4 volumes of ice cold 100% ethanol, incubating overnight at −20°C. Precipitated RNA was gel purified via 7M urea – 8% denaturing polyacrylamide gel electrophoresis (dPAGE) (1x TBE). The RNA was visualized by UV light, excised from the gel, and eluted from the gel by the crush and soak method overnight at 4°C, in ~ 50 mL of diethylpyrocarbonate (DEPC)-treated Milli-Q filtered water. 30K molecular weight cut off (MWCO) Amicon Ultra spin Concentrators (Millipore-Sigma) were used to concentrate the eluted RNA to 2.5 mg/mL, which was then stored at −20°C in DEPC-treated H_2_O until use.

### *In vitro* Xrn1 resistance assay

A total of 5 μg RNA was incubated at 90°C for 3 minutes and 20°C for 5 minutes in 40 μL of refolding buffer (100 mM NaCl, 50 mM Tris pH 7.5, 10 mM MgCl_2_, and 1 mM DTT). Next, 3 μL of recombinant RppH (0.5 μg/μL stock) was added to the mixture which was then aliquoted into two volumes of 20 μL. To one of the aliquots, 1.5 μL of recombinant Xrn1 (0.8 μg/ul stock) was added and the reaction mix was incubated in a thermocycler at 37°C for 2 hrs. 10 μL of RNA from each reaction (+/-Xrn1) was resolved by a 7M urea 8% dPAGE and visualized with ethidium bromide.

### Chemical probing

This method has been adapted from (62). RNA (1.2 pmol) was refolded in 13 μL of folding buffer (77 mM of Na-HEPES pH 8.0) and 4.8 nM of 6-fluorescein amidite-labeled reverse transcription primer (Table S3) at 95°C for 3 min, followed by a 10 min slow cool to room temperature. 2 μL of 100 mM MgCl_2_ were added to the mixture at room temperature. Final concentrations of the folding buffer are as follows: 6.67 mM Na-HEPES pH 8.0, 4.1 nM FAM-labeled RT primers, and 13.3 mM MgCl_2_. This reaction mixture was incubated at 37°C for 15 min. The RNA mixture was centrifuged at maximum speed (30,130 × g) in a tabletop centrifuge, and the full 15 μL were added to one well of the 96 well plate where the remainder of the chemical probing experiments occurred. When setting up this experiment, enough folded RNA was made to fill the contents of a 96 well plate.

Fresh stocks of 0.5% dimethyl sulfate (DMS, Sigma Aldarich, ref: D186309) and 12 mg/mL N-methylisatoic anhydride (NMIA, Invitrogen, ref: M25) were made immediately before probing. Chemical modifier stocks were prepared as follows: 10 μL of DMS was first added to 90 μL of 100% ethanol, which was then added to 900 μL of H_2_O. This 1% DMS solution was then diluted to 0.5% with DEPC treated Milli-q H_2_O. NMIA solution was prepared by dissolving 12 mg of NMIA in dimethyl sulfoxide (DMSO, Acros Organics, ref: 167851000). A stock of DMSO was used as the no modification control. For each condition (DMS, NMIA, or DMSO) 5 μL of the appropriate stock was added to a well containing the 15 μL of folded RNA incubating the reaction at room temperature for 30 min. Specific chemical modifier quenching solutions were prepared which consisted of 1.5 μL of cleaned oligo-dT25 magnetic beads (Invitrogen), 3 μL of 5M NaCl, 0.25 μLof DEPC treated Milli-q H_2_O and 5 μL of the respective quenching solution. Fresh 2-Mercaptoethanol (Sigma, ref: M3148) was used to quench the DMS reaction, 0.5 M 2-(N-Morpholino) ethanesulfonic acid sodium salt (Na-MES, J.T. Baker, ref: 4813-01) to quench NMIA. The appropriate quenching solution was added to each well and incubated at room temperature for 10 min. The chemically modified RNAs bound to the beads were washed with 3x 100 μL of 70% EtOH on a magnetic rack. The washed beads were air dried for ~15 min and then resuspended in 2.5 μLof 2 M betaine (Sigma, ref: B3501).

### Quantitation of chemical probing by reverse transcription

The RNA was reverse transcribed on beads with 2.5 μL of reverse transcription mixture comprised of 1.0 μL of 5x First strand Buffer (Thermo Fisher), 0.25 μL of 0.1 M 1,4-Dithiothreitol (DTT), 0.4 μL of a 10 mM equimolar mixture of dNTPs, 0.75 μL of DEPC-Treated H_2_O, and 0.1 μL of Superscript III reverse transcriptase (Fisher). The mixture was incubated in a 52°C water bath for 50 min. Upon completion, 5 μLof 0.4 M NaOH was added to each well and incubated at 65°C for 20 min to hydrolyze the RNA. The samples were then cooled in an ice bath for 2 min and each reaction was neutralized with 5 μL of an acid quench mixture (1.4 M NaCl, 0.6 N HCl, and 1.3 M NaOAc). The supernatant was aspirated from each well, and the complementary DNA (cDNA) bound to the beads were washed three times with 100 μL of 70% EtOH and allowed to air dry for ~15 min. The cDNA was then eluted off the beads in 11 μL of a ROX-formamide mixture (2.75 uL of ROX-500 ladder (Applied Biosystems) in 1.2 mL of HiDi-formamide (Applied Biosystems), incubating at room temperature for 15 min. The 11 μL cDNA mixture was transferred to an optical plate and analyzed with an ABI 3500 Genetic Analyzer Capillary Electrophoresis machine.

### Preparation of the sanger sequence ladder

Alongside the reverse transcription for the chemically probed RNAs, Sanger sequencing ladders were constructed through reverse transcription of unmodified xrRNA in the presence of dideoxynucleoside triphosphates (ddNTP′s). In brief 2.5 μL of an RNA mixture consisting of 0.2 μL of 0.25 μM FAM primer, 1.5 μL of oligo-dT beads (Invitrogen), 0.5 μL of 2.4 μM RNA, and 0.25 μL of 5 M betaine was added to 2.5 μL of ladder reverse transcription mix consisting of 1.0 μL of 5x First strand Buffer (Thermo Fisher), 0.25 μL of 0.1M DTT, 0.4 μL of 1.0 mM dNTP′s (equimolar dATP, dTTP, dCTP, dGTP), 0.4 μL of 1.0 mM of the appropriate ddNTP, 0.1 μL of Superscript III reverse transcriptase, and 0.25 μL of 5 M Betaine. The outlined reverse transcription protocol was followed as described above.

### Analysis of chemical probing

The HiTRACE MATLAB suite (https://ribokit.github.io/HiTRACE/) with MATLAB (v8.3.0.532) was used to analyze the chemical probing data as described in (63). CE traces were first aligned manually and verified using the Sanger sequencing ladder in HiTRACE. Each run was corrected for signal attenuation, background subtracted with values from the DMSO channels, and then normalized via the 5′ and 3′ hairpin loops (GAGUA) flanking the sequence of interest in the construct. The nucleotide reactivities for each modifier were calculated, and the reactivities were mapped back to the secondary structure of the xrRNA using Varna v. 3.93 (64).

## ACKNOWLEDGEMENTS

The authors thank Dr. Esteban Finol for sharing sets of relevant sequence data, Dr. Elena Rivas for her help with R-scape, and Drs. Rene Olsthoorn and Peter Bredenbeek for useful communication and insight regarding the presence of xrRNAs in Pegiviruses. We also thank David Farrell for computational assistance, Dr. Anna-Lena Steckelberg and David Costantino for critical reading of the manuscript and insightful comments, as well as current and former Kieft Lab members for thoughtful discussions and technical assistance. This work was supported by NIH grants R35GM118070 and R01AI133348 (JSK), and T32AI052066 (MJS).

## Supplemental Material

**Table S1:**
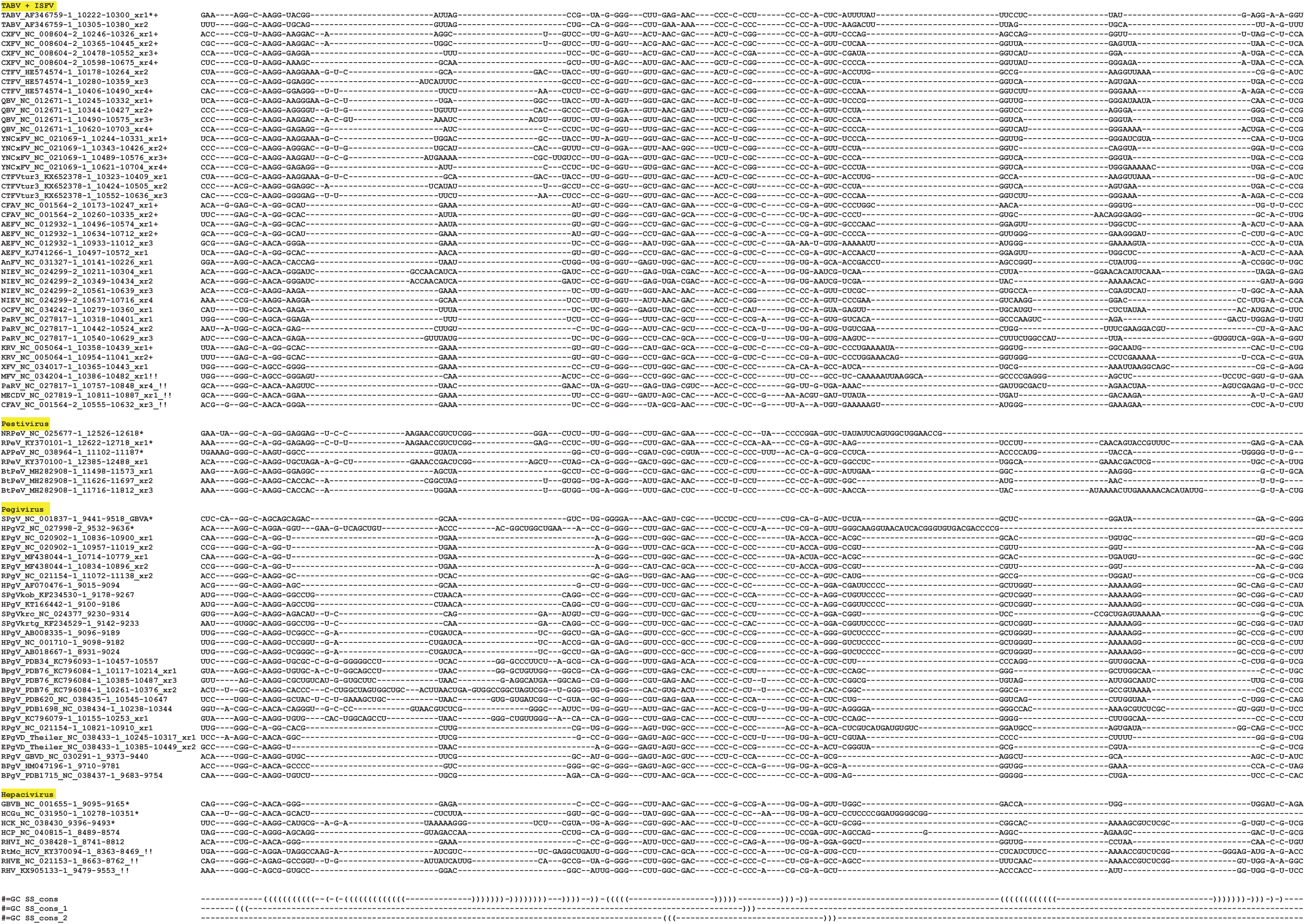
Complete subclass 1b xrRNA sequence alignment. This sequence alignment was used to generate the subclass 1b covariance model (Fig. 2B). Constructs used for chemical probing analysis are designated by a star. Labels are formatted with the name of the virus, the accession number, the location of the xrRNA within the accession number, and the relative position of the xrRNA, when several are found in series (i.e., xr1 is the most 5′ xrRNA element). Sequences ending in a “+” were used in the initial sequence alignment as input for *Infernal* searches.

**Table S2:**
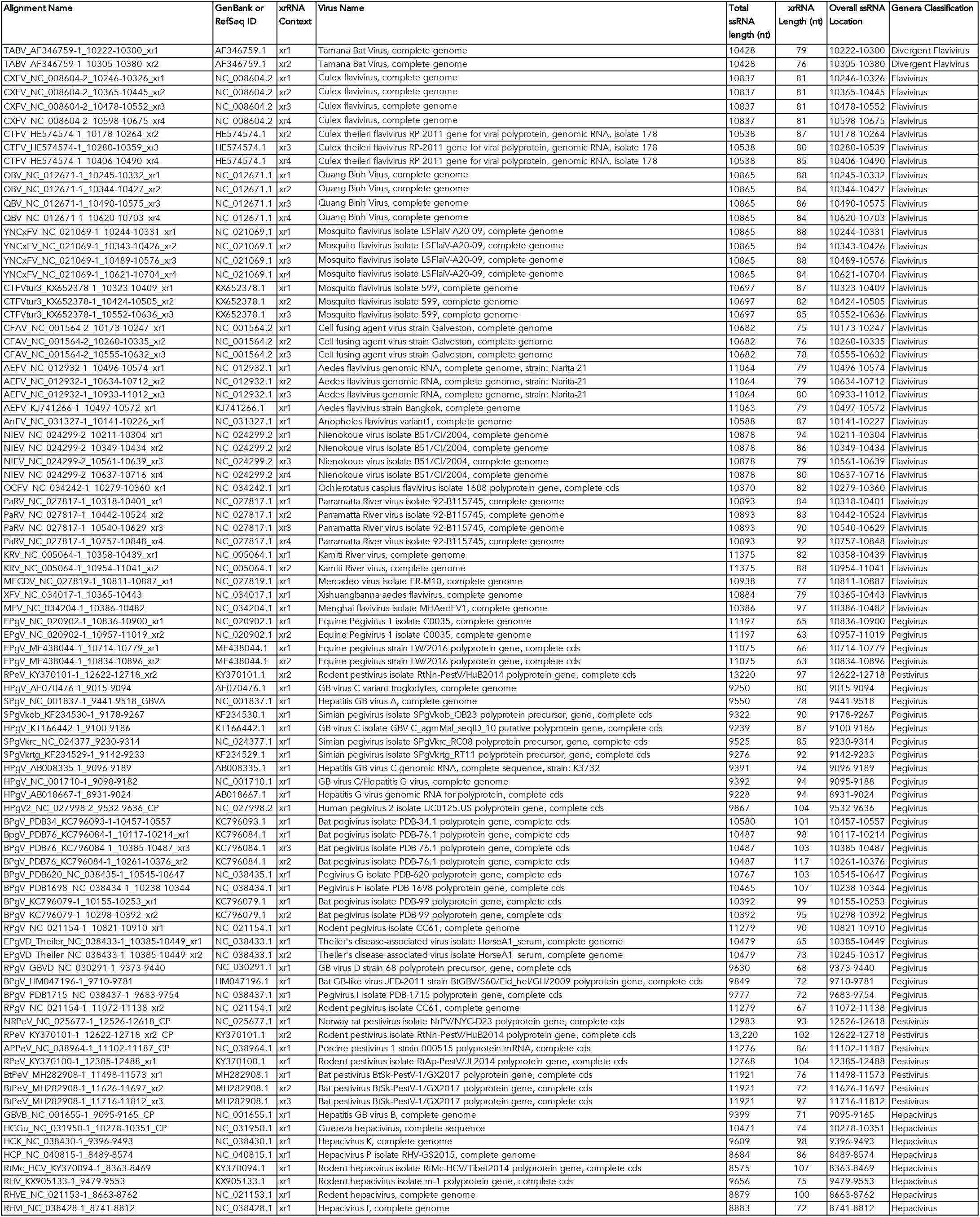
General information about subclass 1b xrRNA sequences.

**Table S3:**
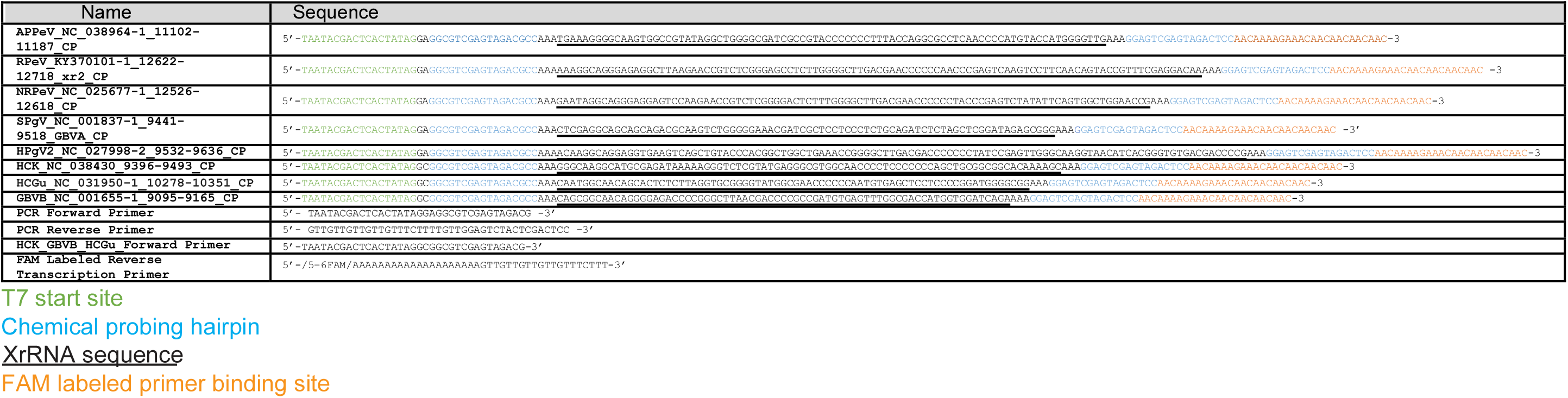
Subclass 1 xrRNA construct and primer information. Sequence information for the biochemically tested subclass 1b xrRNA, and PCR and reverse transcription primers used for cloning and reverse transcription.

**Figure S1:**
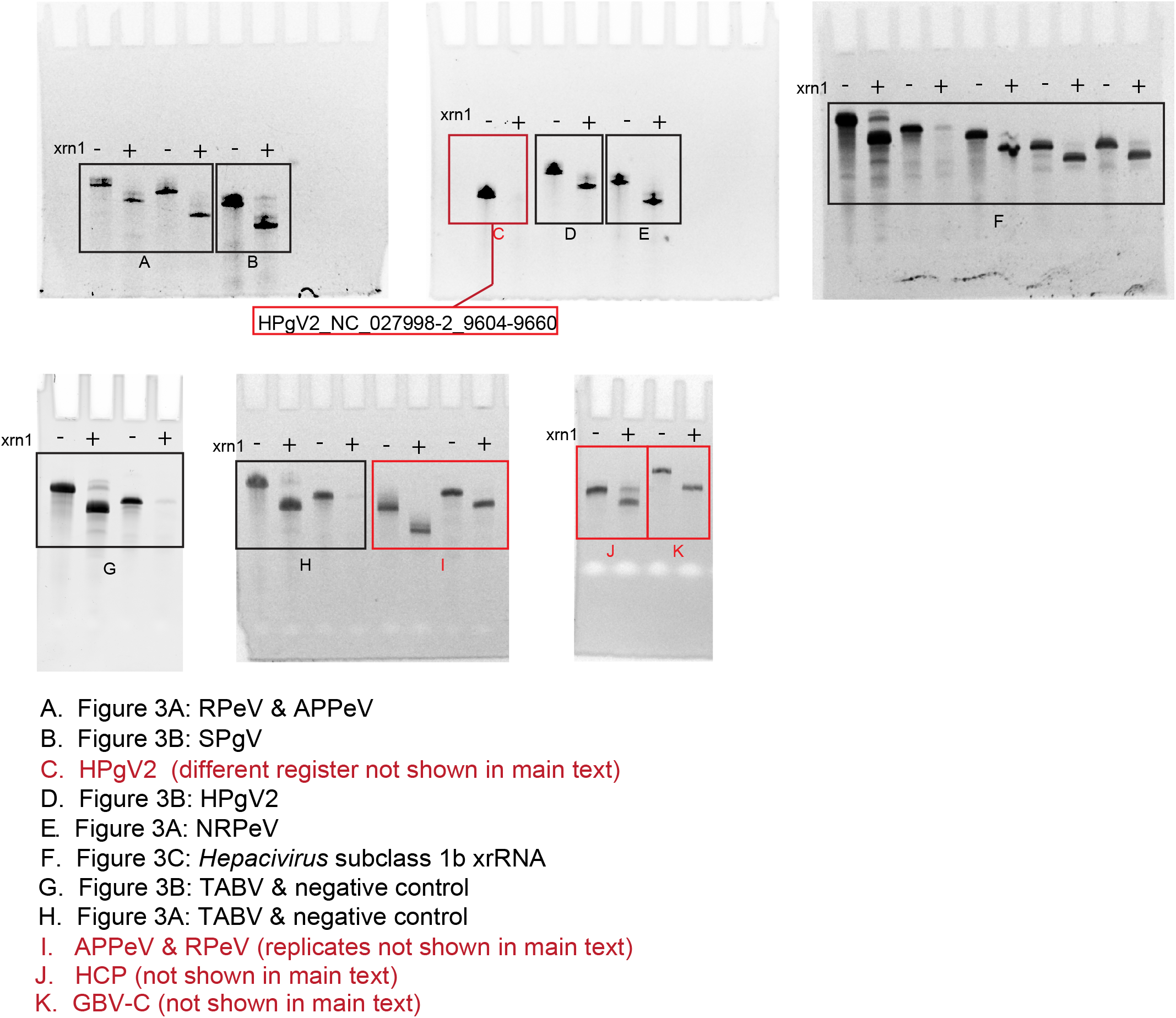
Uncropped subclass 1b *in vitro* degradation assay gels. Raw gel images used and cropped for Figure 3. Red boxes indicate data only shown in the supplementary information and not in the main text.

**Figure S2:**
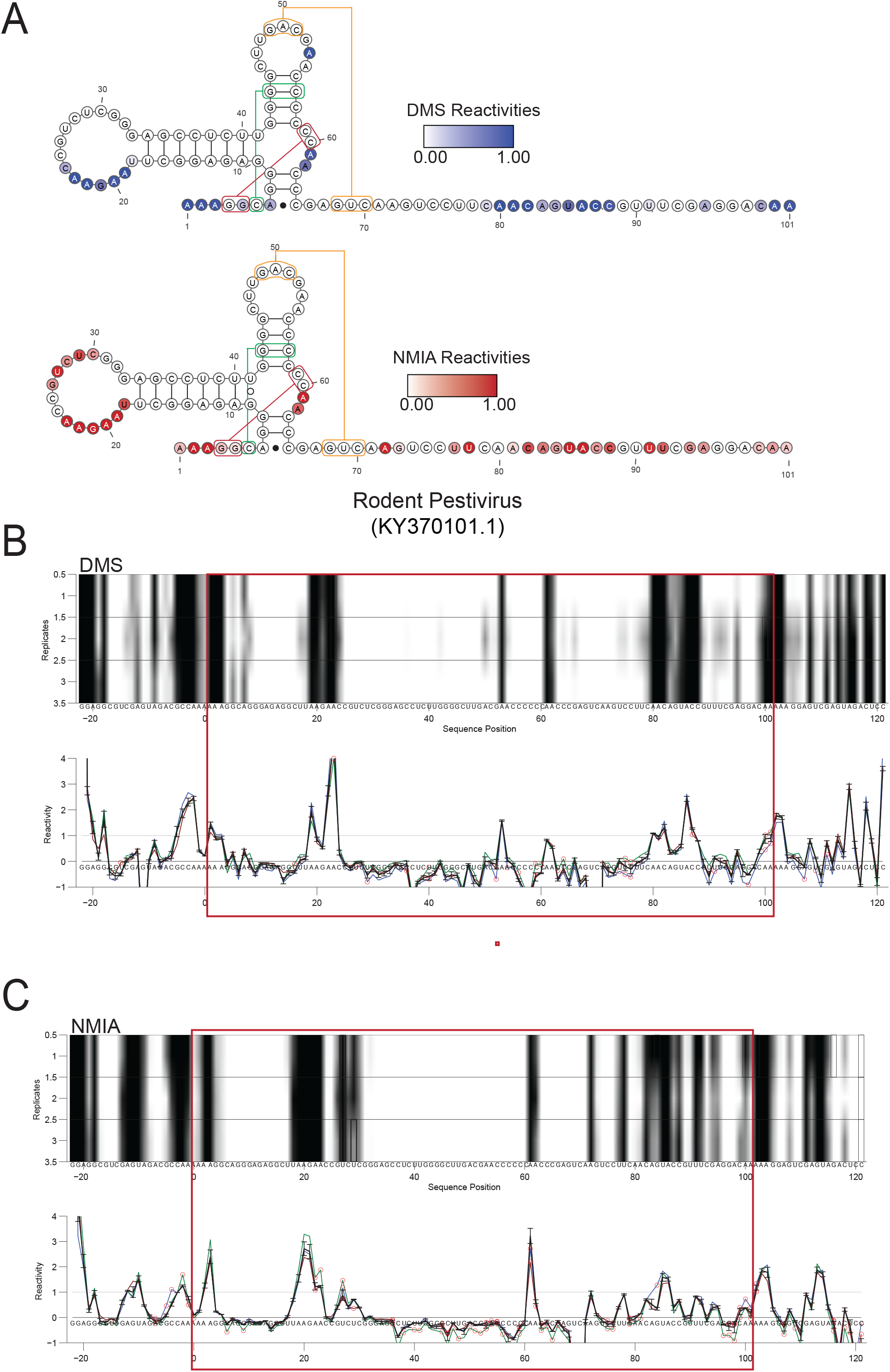
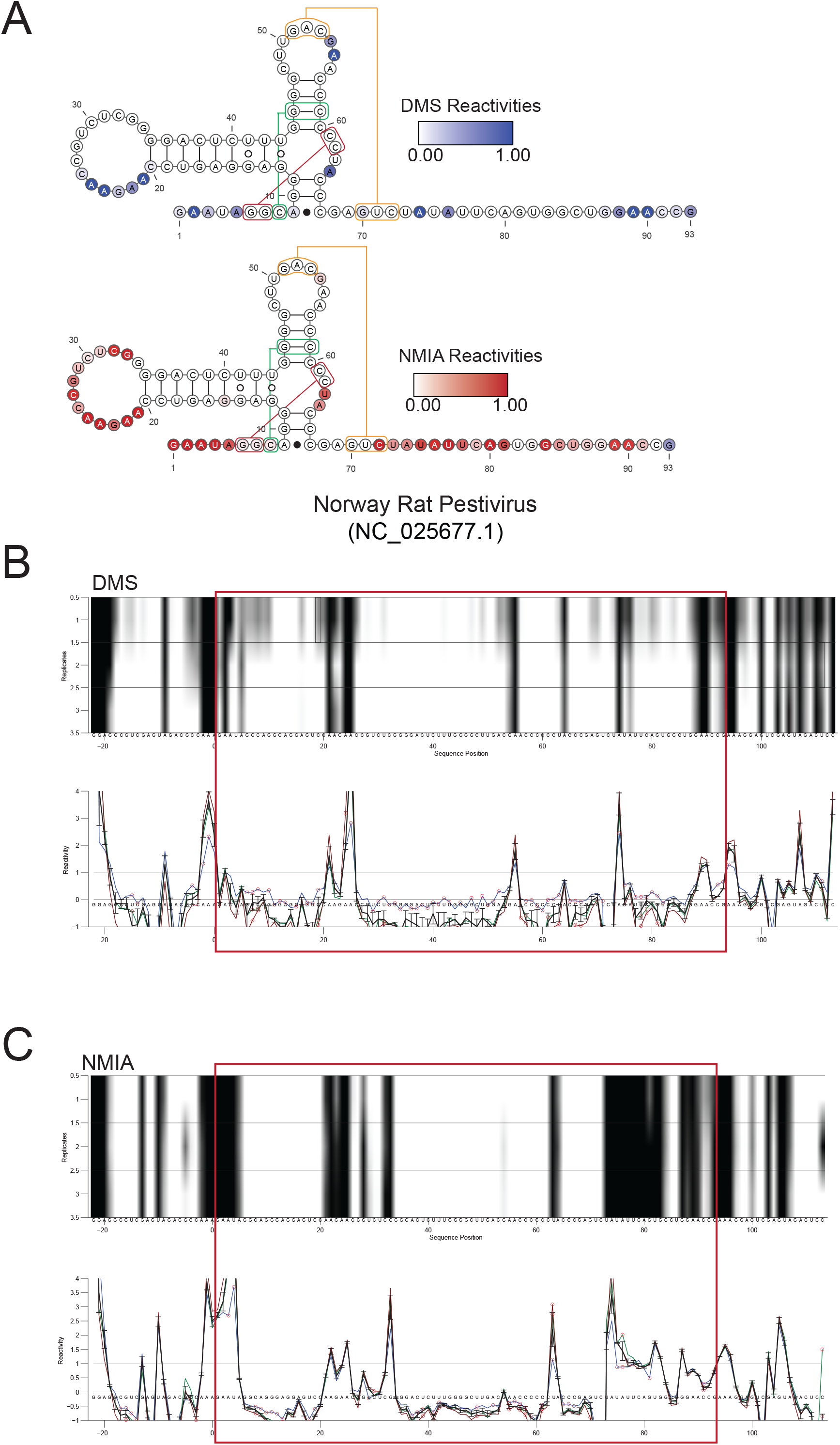
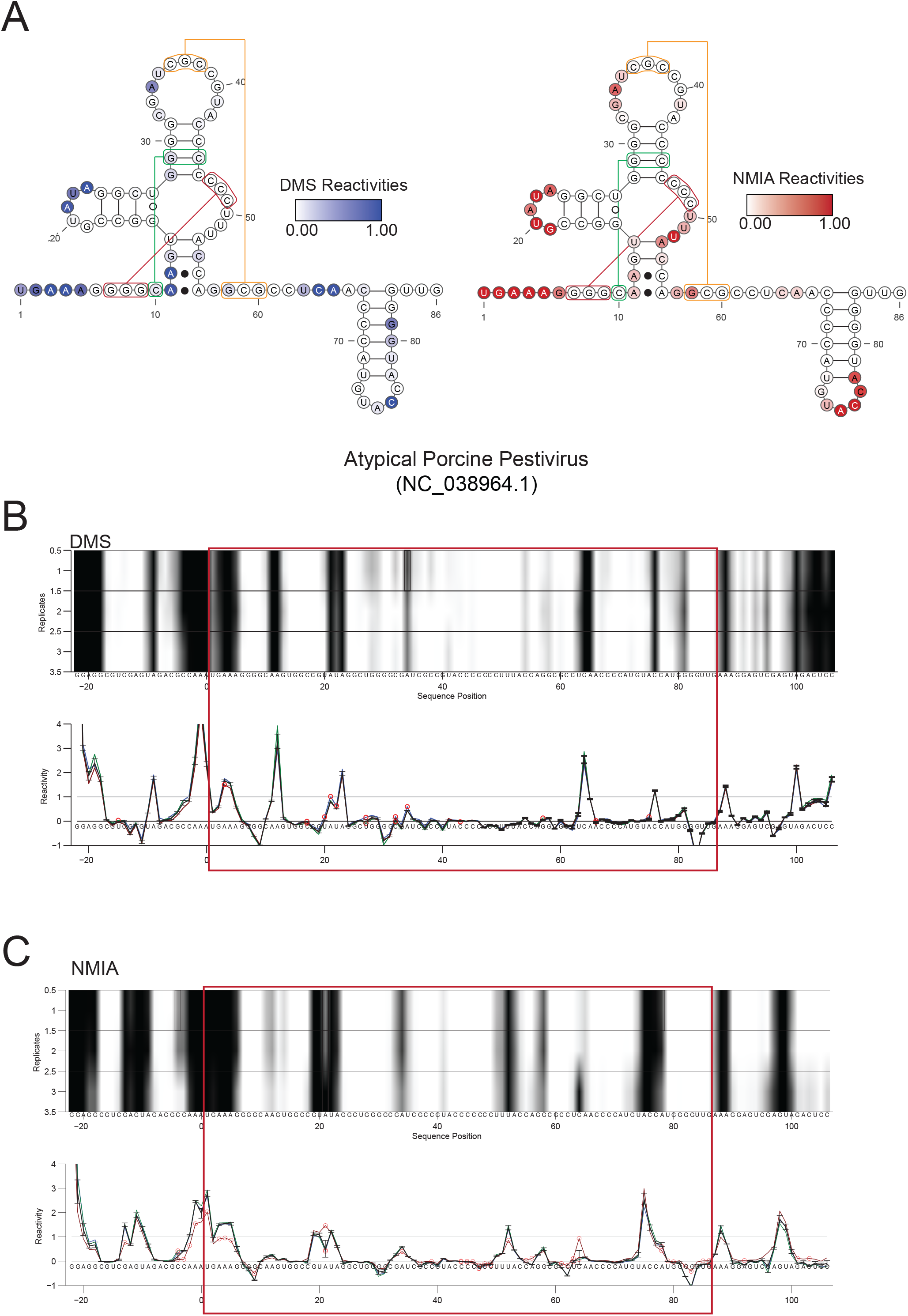
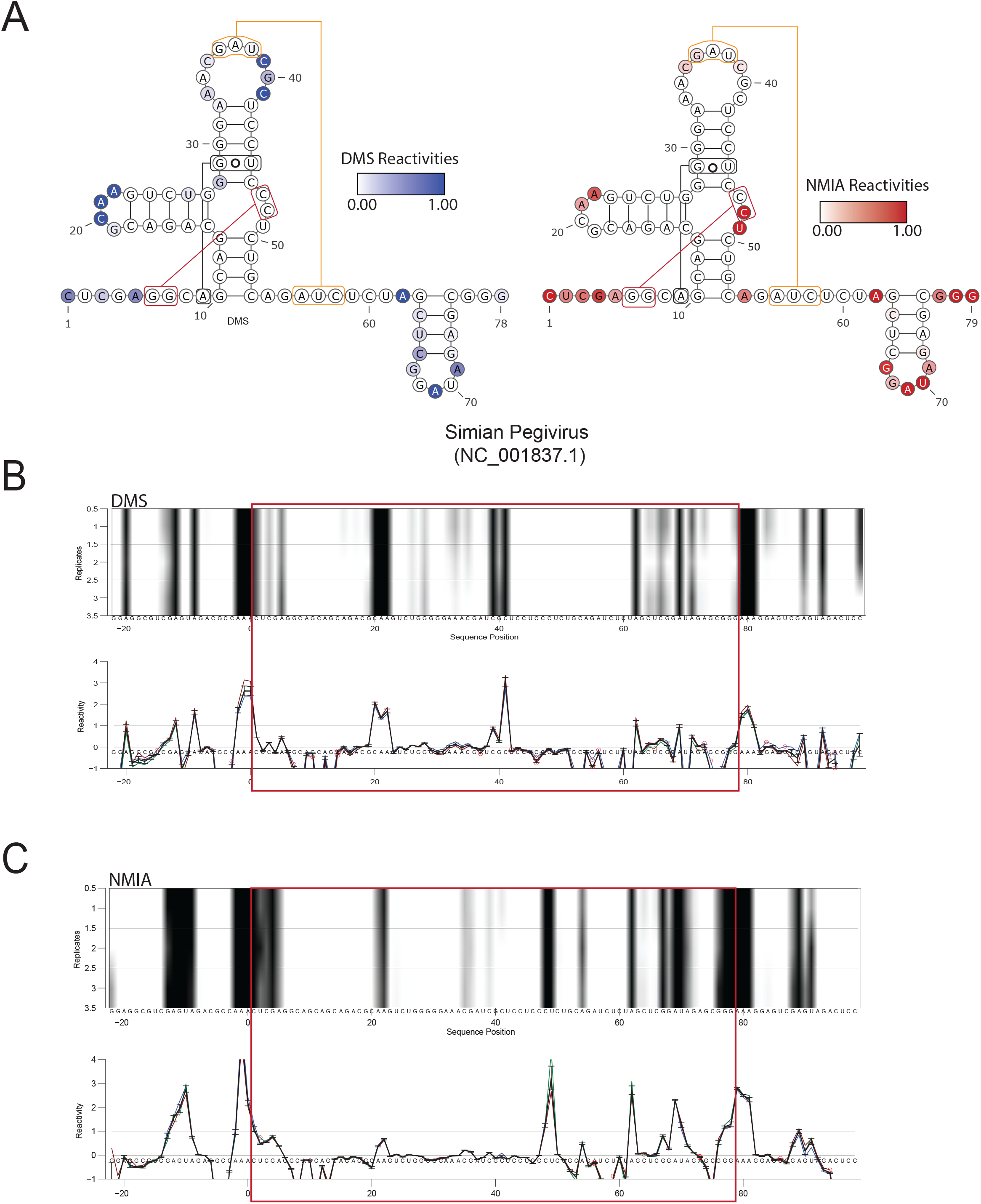
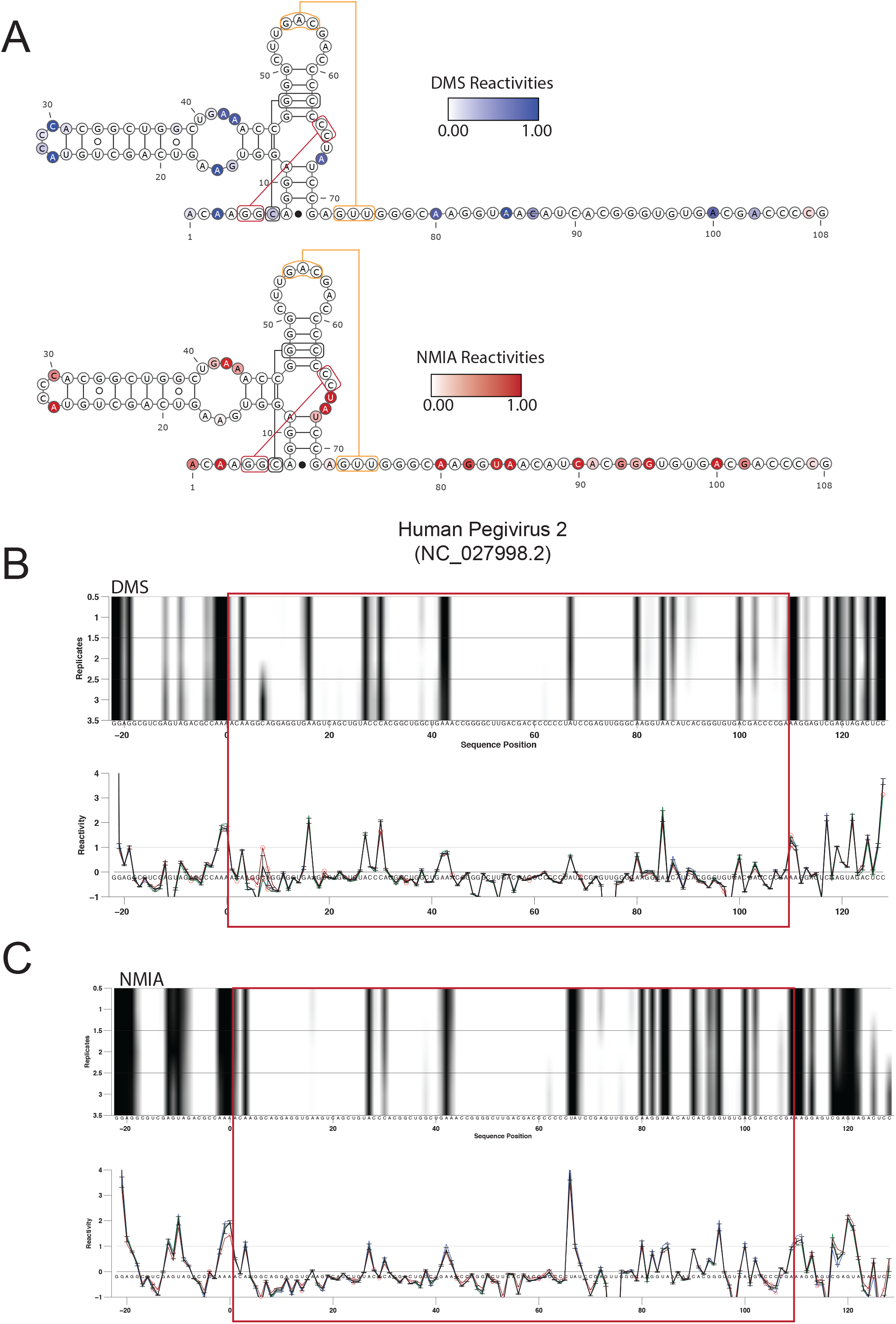
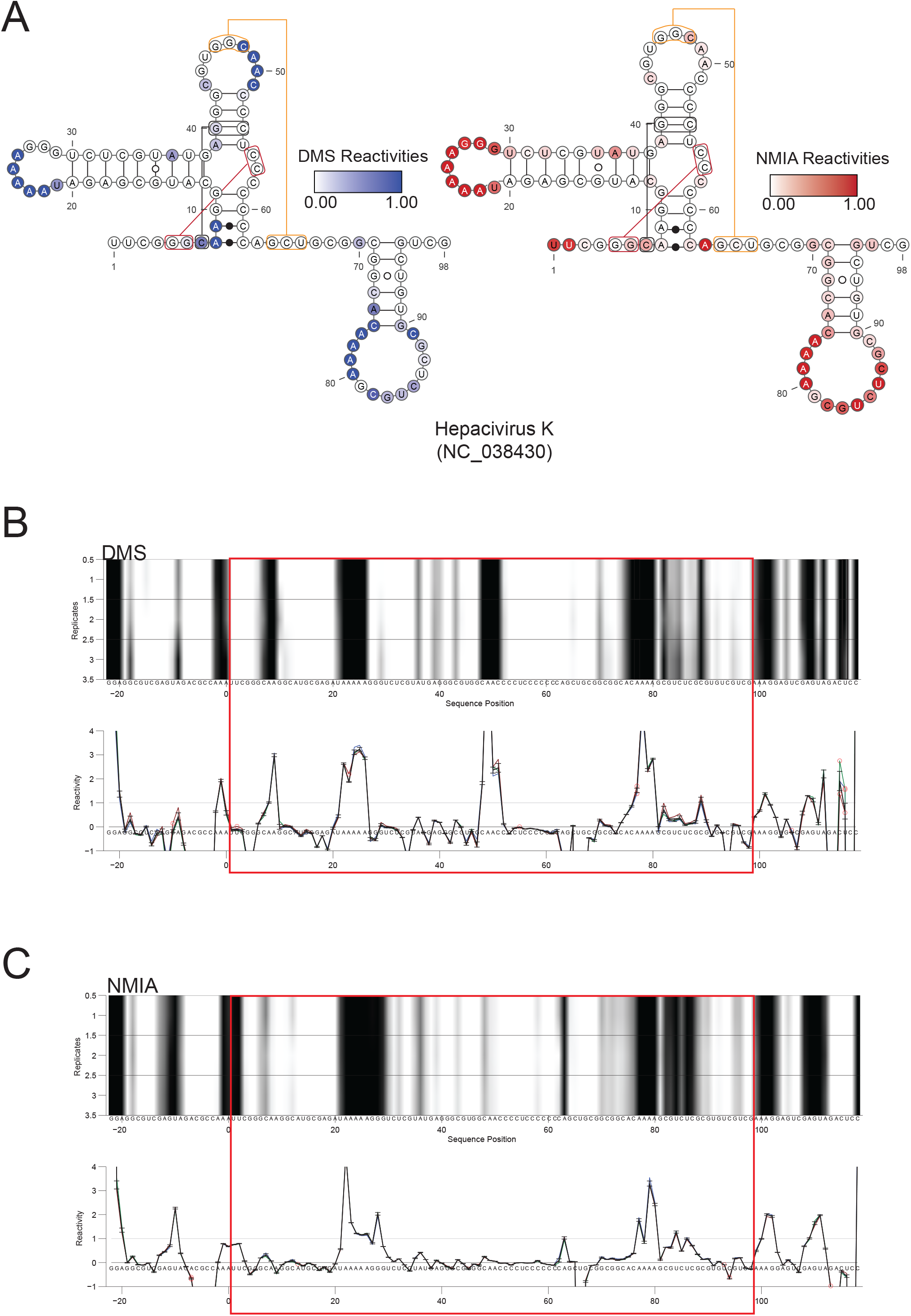
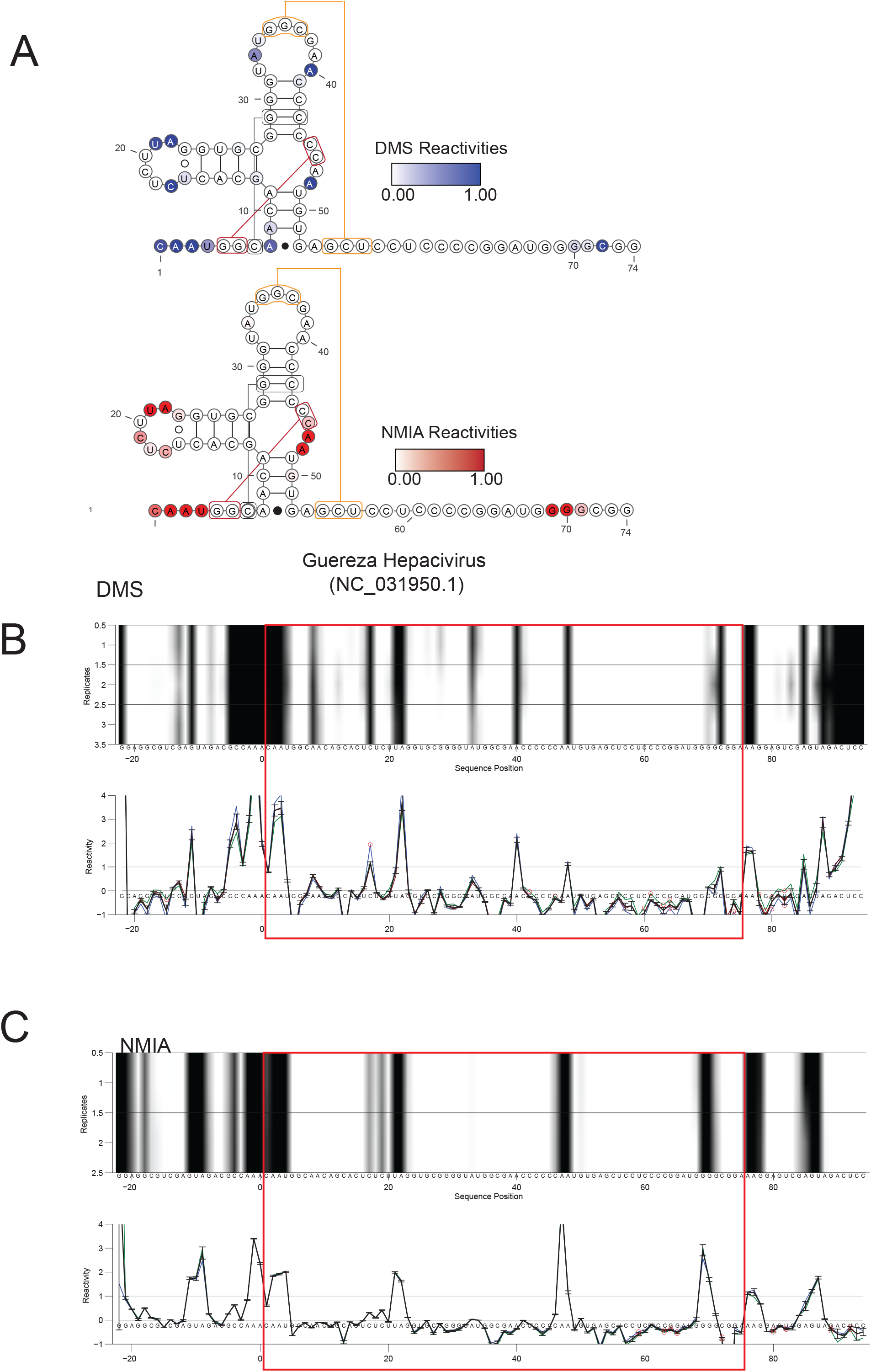
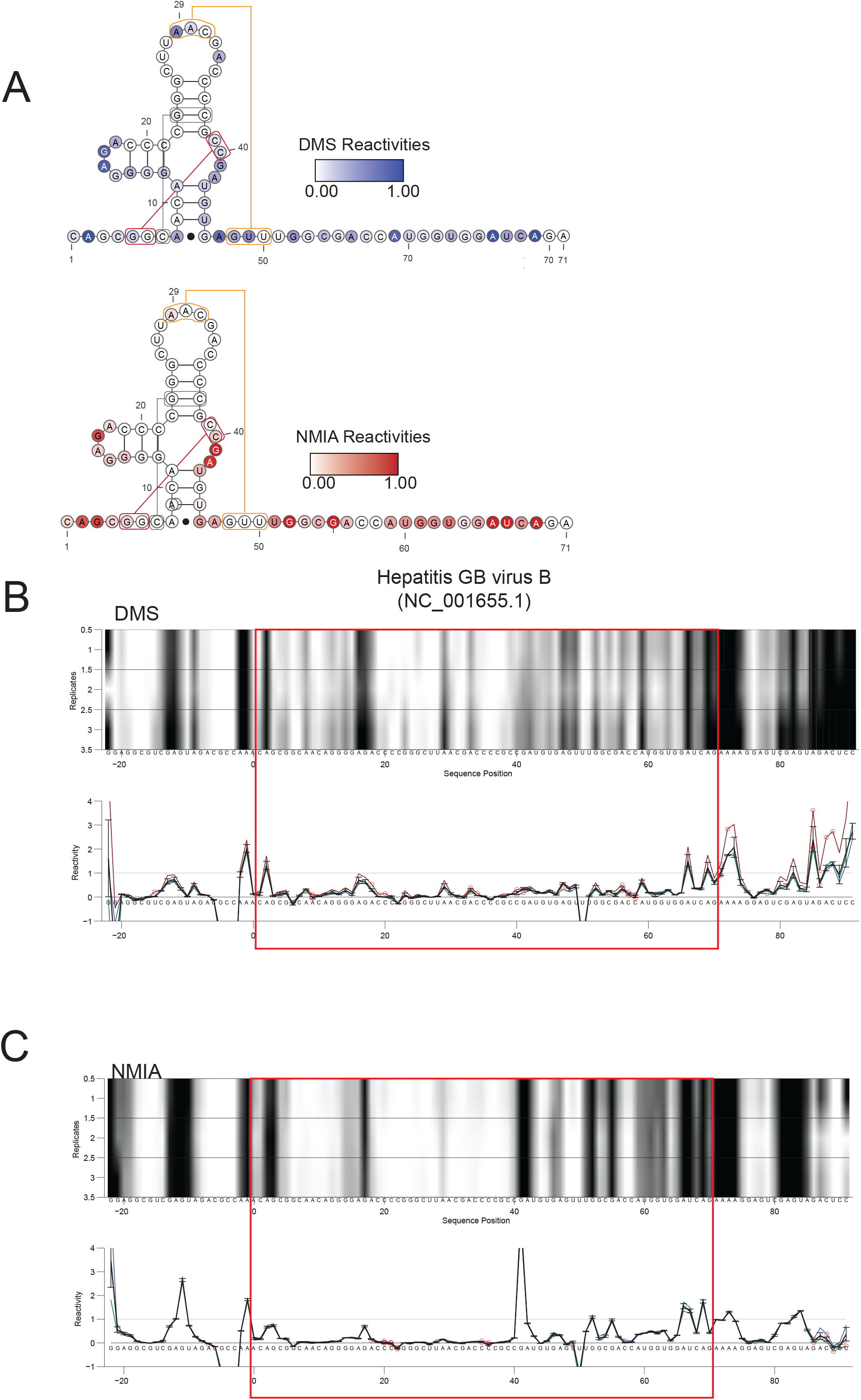
Secondary structure validation of subclass 1b xrRNA by chemical probing. Each of the following pages contains a single xrRNA construct with similar labels and notation, showing the raw data for chemical probing experiments. The eight constructs are ordered by genera: *Pestivirus, Pegivirus, and Hepacivirus.* (A) Normalized DMS and NMIA reactivities mapped onto the secondary structure representation of the xrRNA, including the putative P4 region. (B,C) CE lanes and reactivity plot for DMS (B) and NMIA (C) used in the experiment. Red circles around the peaks in the reactivity plot represent a higher degree of standard deviation observed for that point.

## Notes

### Competing Interest Statement

The authors have declared no competing interest.

